# Karyopherin Kap114p regulates TATA-binding protein-mediated transcription

**DOI:** 10.1101/411223

**Authors:** Chung-Chi Liao, Sahana Shankar, Golam Rizvee Ahmed, Kuo-Chiang Hsia

**Affiliations:** Institute of Molecular Biology, Academia Sinica, Taipei 11529, Taiwan; Institute of Biochemistry and Molecular Biology, College of Life Sciences, National Yang-Ming University, Taipei 11221, Taiwan

## Abstract

Gene expression is regulated by nuclear accessibility of transcription factors controlled by transport receptors and competitive inhibitors. Multiple karyopherin-βs (Kap-βs) facilitate nuclear import of yeast TATA-binding protein (yTBP). However, only Kap114p suppresses temperature-sensitive yTBP mutations, suggesting Kap114p executes undefined non-transport functions. We show that yTBP preferably binds Kap114p with an affinity three orders of magnitude higher than other Kap-βs facilitating yTBP nuclear import. Our crystal structure of Kap114p reveals an extensively negatively-charged concave surface, accounting for high-affinity basic-protein binding. Furthermore, we biochemically demonstrate that two intra-HEAT-repeat inserts act as regulatory TBP-binding domains carried by TBP-associated factor 1 (TAF1-TAND), suppressing binding of yTBP with DNA and the transcription factor IIA. Remarkably, dual-knockout of *Kap114* and *TAF1-TAND* in yeast synergistically suppresses cell growth. Our study reveals that Kap114p has a dual function that modulates the nuclear localization and activity of yTBP, illuminating how the nuclear transport machinery regulates yTBP-mediated transcription.

## INTRODUCTION

In eukaryotes, the nuclear envelope (NE) separates nuclear transcription and cytoplasmic translation. Transcription factors synthesized in the cytoplasm need to be imported into the nucleus, whereas the translational machinery (e.g. ribosomal subunits) assembled in the nucleus has to be exported to the cytoplasm. A family of soluble transport factors, termed karyopherin-β (Kap-β) in yeast, facilitates the majority of nucleo-cytoplasmic transport. Kap-βs recognize nuclear localization signals (NLS) and nuclear export signals (NES) carried by cargo molecules (either directly or through an adapter, karyopherin-α) to transfer cargoes from one side of the NE-embedded nuclear pore complex (NPC) to the other. Assembly of the cargo• Kap-β complex is modulated by an asymmetric distribution of RanGTP across the NE. A high nuclear concentration of RanGTP unloads imported cargoes from Kap-βs and promotes complex assembly of exported cargoes and Kap-βs, whereas a low cytoplasmic RanGTP level releases exported cargoes from Kap-βs and allows formation of the imported cargo•Kap-β complex.

Nuclear transcription is performed by three distinct RNA polymerases—RNA polymerase I, II and III (Pol I, II and III)—that are all mediated by an essential, evolutionarily-conserved TATA-binding protein, TBP ^1^. During RNA Pol II-mediated transcription, TBP recognizes the TATA box sequence of the promoter and assembles the pre-initiation complex (PIC) by recruiting other proteins (e.g. transcription factor IIA, TFIIA) ^2^. In current models, the nuclear function of TBP is modulated by two groups of proteins: (1) nuclear transport factors, with self-dimerization of TBP increasing its molecular weight beyond the diffusion limit (∼40 kDa) of the NPC and thereby preventing its nuclear localization ^3-5^; and (2) TBP-associated factors (TAFs) that produce a competitive inhibition network by forming heterocomplexes with TBP ^6^. Notably, multiple Kap-βs have been shown to mediate nuclear import of yeast TBP (yTBP) (e.g. Kap95p, Kap114p, Kap121p and Kap123p) ^7,8^. However, suppression of temperature sensitivity caused by yTBP mutations within the DNA-binding pocket (Y94C) or TFIIA interaction site (K133 and 138L) was specifically attributable to Kap114p ^7^. These results led us to hypothesize that the nuclear transport factor Kap114p carries out functions other than nuclear import of yTBP and thus can rescue the yTBP mutant phenotypes.

To prove our hypothesis, we first show that yTBP binds to Kap114p with an affinity that is three orders of magnitude higher than for other examined Kap-βs (Kap95p and Kap121p). Next, our crystal structure of Kap114p reveals that Kap114p structurally resembles a cargo-bound form of exportin Cse1p, whereas the highly negative surface charges of Kap114p allows it to bind nucleic acid-binding proteins, such as yTBP. Biophysical and biochemical analyses together with our crystal structure information revealed two inserts within HEAT repeats (HEAT8 hairpin and HEAT19 loop) bearing protein sequence compositions and secondary structure elements similar to the N-terminal domain of TAF1 (TAF1-TAND) that modulates binding of yTBP to DNA and TFIIA. Moreover, yeast genetic analyses revealed that knockout of *Kap114* and *TAF1-TAND* has a synergistic effect on cell growth. Hence, apart from conducting nuclear import like closely-related Kap95p and Kap121p, Kap114p acts as a negative TBP-associated factor (e.g. TAF1) that suppresses yTBP activities in the nucleus.

## RESULTS

### Biochemical characterization of yTBP and Kap-βs interaction

yTBP localization to the nucleus is facilitated by different Kap-βs, such as Kap95p, Kap114p and Kap121p ^7,8^. To biochemically validate the many interactions that occur between yTBP and different Kap-βs, we first conducted pull-down assays using purified recombinant proteins. Interestingly, while constitutively-active His-tagged RanQ69L pulled down comparable amounts of Kap95p, Kap114p and Kap121p, pull-down of Kap114p by GST-fused yTBP (aa 61-240; hereafter yTBP) was substantially elevated compared to that of Kap95p and Kap121p (Fig. 1a,b), suggesting that yTBP directly binds Kap114p with a relatively higher binding affinity. Next, we used isothermal titration calorimetry (ITC) to measure dissociation constants (Kds) between yTBP and Kap-βs. The Kd values of yTBP and Kap95p or Kap121p were ∼ 10 and 15 μM, respectively (Fig. 1d,e). Notably, under the same experimental conditions, ITC revealed the binding affinity of Kap114p to yTBP (6 nM) to be three orders of magnitude higher than for Kap95p and Kap121p (Fig. 1c), further demonstrating that yTBP preferably binds Kap114p. Additionally, the respective biphasic binding profile ITC revealed that multiple binding sites contribute to the interaction between yTBP and Kap114p (Fig. 1c).

**Fig. 1:**
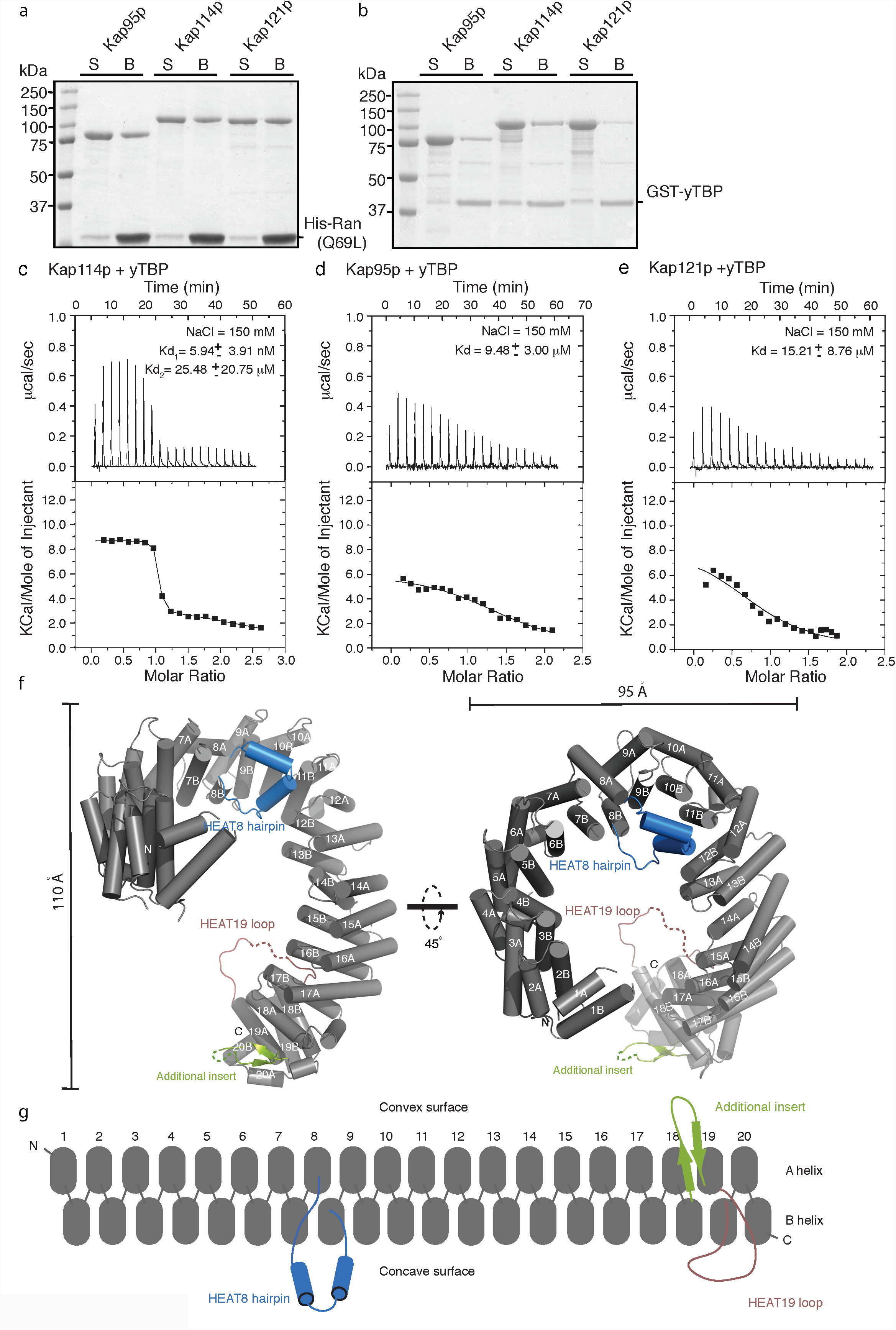
Biochemical and Structural characterization of yTBP and Kap114p interaction. **a,b,** Pull-down assays of RanQ69L and yTBP (61-240) with Kap95p, Kap114p and Kap121p. **(a)** His-tagged RanQ69L and **(b)** GST-fused yTBP (61-240) were incubated with recombinant Kap95p, Kap114p and Kap121p. Unbound (S) and bound (B) samples were analyzed by SDS-PAGE and stained with Coomassie blue. **c,d,e,** ITC titration curves (upper) and binding isotherms (lower) of **(c)** Kap114p, **(d)** Kap95p, and **(e)** Kap121p with yTBP (61-240). Salt concentration and Kd values are indicated. **(f)** Cartoon representation of the Kap114p crystal structure. The A and B helices composed of each HEAT repeat are shown as gray cylinders. The HEAT8 hairpin, HEAT19 loop and HEAT18/19 loop are highlighted in blue, red and green, respectively. A 45° rotated view is shown on the right. Overall dimensions of the Kap114p structure are indicated. **(g)** Schematic representation of the Kap114p domain configuration using the same color code as shown in **f**.

### The Kap114p crystal structure shows an exportin Cse1p-like fold

To gain structural insights into how Kap114p binds to yTBP with an affinity in the sub-nanomolar range, we determined the crystal structure of Kap114p. Crystallization of Kap114p in citric acid, bis-tris propane (pH 5.8), PEG3350, EDTA, and ethanol generated hexagonal crystals (space group P3_1_2_1_), with one molecule per asymmetric unit. Phase determination was carried out using data collected from seleno-L-methionine-labeled crystals at selenium peak and inflection wavelengths (Supplementary Table 1). The final structure was refined to a resolution of 2.5 Å with an *R*-factor of 21.1% (*R*_free_ 26.4%), and there were no outliers in the Ramachandran plot (Supplementary Table 1). Kap114p comprises 20 HEAT repeats that form a right-handed superhelix structure (Fig. 1f,g). Additional insertions of a helix-turn-helix hairpin and two loops are found in the HEAT8, HEAT19 and HEAT18/19 repeats, respectively (Fig. 1f,g). Superimposition of Kap114p with the crystal structure of the cargo-bound exportin Cse1p illustrates striking structural similarities between their tertiary structures (Supplementary Fig. 1a,b) ^9^. Two intra-HEAT-repeat inserts found in Kap114p, i.e. the HEAT8 hairpin and the HEAT19 loop, also occur in Cse1p (Supplementary Fig. 1b). As Cse1p forms a nuclear export complex together with its cargo and Ran ^9^, we next performed a GST pull-down assay to test if Kap114p spontaneously binds Ran and yTBP (cargo). In the presence of RanQ69L, pull-down of Kap114p by GST-yTBP was substantially reduced compared to pull-down in the absence of Ran (Supplementary Fig. 1c), suggesting the existence of a mutually exclusive Ran and yTBP binding site in Kap114p. Thus, although Kap114p (importin) and Cse1p (exportin) show high structural similarity, Kap114p functions as an import receptor.

Cse1p is an exportin dedicated to nuclear export of yeast Importin-α (Kap60p; isoelectric point (pI)=4.8) ^10,11^, whereas Kap114p imports multiple nucleic acid-binding proteins that bear positively-charged surface patches (e.g. yTBP, histone H2A/H2B and NAP) ^7,8,12,13^. Thus, Kap114p and Cse1p should exhibit different surface charge distributions in order to accommodate binding of their cargoes. Electrostatic surface calculations revealed a neutral charge distribution in the concave surface of the Cse1p central region that is crucial for Kap60p binding (Supplementary Fig. 1d). Interestingly, calculations of electrostatic properties and residue conservation indicated that the concave surface of Kap114p, i.e. where cargoes are supposed to bind, is negatively charged and well conserved (Supplementary Fig. 1e,f).

The crystal structure of cargo-free Kap114p showed an open conformation different from that of Cse1p in the cargo-free state (Supplementary Fig. 1b). We wondered whether this open form of Kap114p can exist in a non-crystalline environment, so we carried out small angle X-ray scattering (SAXS) analyses to obtain information on its global shape in solution. The *P*(*r*) function derived from SAXS data, which reflects the distribution of mean electron density of the molecule, displayed a bimodal distribution with two maxima, implying Kap114p molecule flexibility (Supplementary Fig. 2a). Furthermore, the profile of the Kap114p Kratky plot is comparable to that of CRM1 (Supplementary Fig. 2c), an exportin that adopts different conformations in solution ^14^. This result suggests that open and closed conformations of Kap114p may co-exist in solution. The theoretical scattering profile generated by our Kap114p model exhibited reasonable agreement with our experimental data (Supplementary Fig. 2b). Furthermore, we used the *ab initio* modeling method GASBOR for shape determination, which revealed an open ring-liked conformation that allowed the crystal structure of cargo-free Kap114p to be docked into the SAXS-derived envelope (Supplementary Fig. 2d,e).

**Fig. 2:**
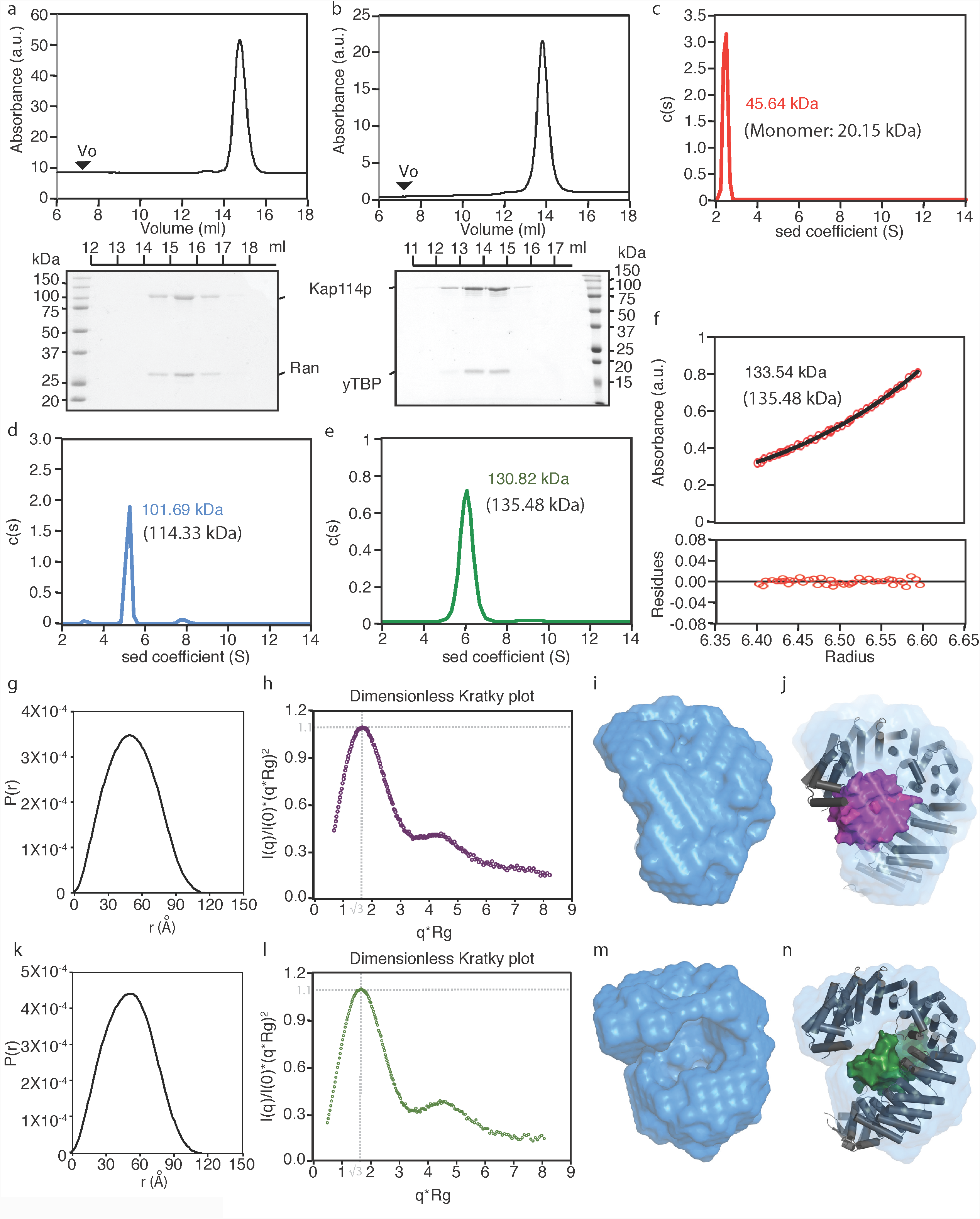
Ran and yTBP binding by Kap114p. **a,b,** Purified recombinant Kap114p was mixed with RanQ69L or yTBP and anaylzed by SEC. SEC (Superdex 200) elution profiles of the **(a)** Kap114p•RanQ69L and **(b)** Kap114p•yTBP complexes. Peak fractions were analyzed by SDS-PAGE and stained with Coomassie blue. The void volume (*Vo*) of the peak fraction and absorbance (a.u.) at 280 nm for each complex is indicated. **c,d,e,** SV-AUC analysis of **(c)** yTBP, **(d)** Kap114p and the **(e)** Kap114p•yTBP complex. The experimental data were analyzed by the Sedfit program and are presented above molecular weights calculated using standards (in brackets). **f,** SE-AUC analysis of the Kap114p•yTBP complex. The average molecular mass determined by the Sedfit program was about 133.5 kDa. The lower panel shows the residual difference (standard deviation) between experimental and fitted values. **g,k,** Plots of the distance distribution function P(r) for the **(g)** Kap114p.RanQ69L and **(k)** Kap114p•yTBP complexes derived from experimental SAXS data using the GNOM program. **h,l,** Rg-based dimensionless Kratky plot of the Kap114p.RanQ69L **(h)** and Kap114p•yTBP **(l)** complexes. Intersection of the two dotted gray lines indicates features of a compact protein. The *ab initio* envelopes determined by GASBOR for the **(i)** Kap114p•RanQ69L and **(m)** Kap114p•yTBP complexes. Models of the **(j)** Kap114p.RanQ69L and **(n)** Kap114p.yTBP complexes derived from SAXS profiles using FoXSDock and docked into the *ab initio* envelopes obtained from GASBOR.

### Kap114p interacts with Ran and yTBP with equal stoichiometry

We next examined in molecular detail how Kap114p interacts with Ran and yTBP. We biochemically reconstituted Kap114p•RanQ69L and Kap114p•yTBP complexes using recombinant Kap114p, RanQ69L and yTBP. Purified Kap114p formed stable complexes with either RanQ69L or yTBP, co-migrating with either one in a sharp mono-dispersed peak in size exclusion chromatography (SEC; Fig. 2a,b). The band intensities of Kap114p and Ran on SDS-PAGE suggest a 1:1 equal stoichiometry (Fig. 2a), consistent with the ratios found in all Ran•Kap-β complexes ^15^. However, since yTBP has been reported to form dimers in solution ^3^, the stoichiometry of the Kap114p•yTBP complex needed further assessment. Thus, we applied analytic ultracentrifugation with sedimentation velocity (SV-AUC) and sedimentation equilibrium (SE-AUC) to characterize the size distributions of yTBP, Kap114p and the Kap114p•yTBP complex. SV-AUC revealed that yTBP and Kap114p alone are predominantly a homo-dimer and monomer, respectively, in solution, with molecular masses consistent with calculated values (Fig. 2c,d). Sedimentation profiles from SV-AUC and SE-AUC of the Kap114p.yTBP complex revealed a molecular mass of ∼130 kDa (Fig. 2e,f), concordant with the calculated molecular weight of a hetero-dimeric complex, suggesting a 1:1 stoichiometry for the Kap114p.yTBP complex.

Next, we carried out SAXS analysis to determine the global shapes of Kap114p in complex with either RanQ69L or yTBP. The *P*(*r*) function derived from SAXS data showed a single peak for both complexes, indicating a single conformation in solution and differing from our assessment of Kap114p alone (Fig. 2g,k and Supplementary Fig. 2a). The bell-shaped Kratky plots of both complexes had maxima of ∼1.1 at q* q*Rg=√3, further suggesting a compact folded conformation (Fig. 2h,l). Next, we applied GASBOR *ab initio* modeling to generate 10 independent and reproducible models—Normalized spatial discrepancy (NSD)=1.29 ± 0.03 for Kap114p•yTBP and NSD=1.31 ± 0.06 for Kap114p•RanQ69L—that matched well with our experimental data (Supplementary Fig. 2f,h). The final SAXS envelopes of both complexes displayed a compact globular shape without a central cavity (Fig. 2i,m). These results demonstrate that Ran and yTBP associate with Kap114p, filling the central cavity of the ring-like structure. To dock crystal structures into the SAXS-derived envelopes, we used FoXSDock that predicts interfaces based on experimental data and calculated energies to best fit the SAXS profiles. FoXSDock could place either RanQ69L or yTBP in the center of the horseshoe-shaped Kap114p structure, with the two arms of the horseshoe wrapped around either protein (Fig. 2j,n and Supplementary Fig. 2g,i).

### The HEAT8 hairpin of Kap114p facilitates Ran-binding

Our low-resolution models showed that the HEAT8 hairpin and HEAT19 loop facing the concave surface of Kap114p might physically contact Ran and yTBP. To examine their Ran-and yTBP-binding activities, we biochemically purified two deletion constructs lacking, respectively, the HEAT8 hairpin (amino acids 347-371, hereafter Kap114p (Δ347-371)) and the HEAT19 loop (aa 899-956, hereafter Kap114p (Δ899-956)) using multiple steps of a chromatographic approach and further assessed both constructs by mass spectrometry, SEC, and circular dichroism (CD) (Fig. 3a, b and Supplementary Fig. 3a-e). The protein secondary structure composition and solution properties of wild-type Kap114p and the two deletion mutants were comparable (Supplementary Fig. 3a-e), so they were suitable for further biochemical studies.

**Fig. 3:**
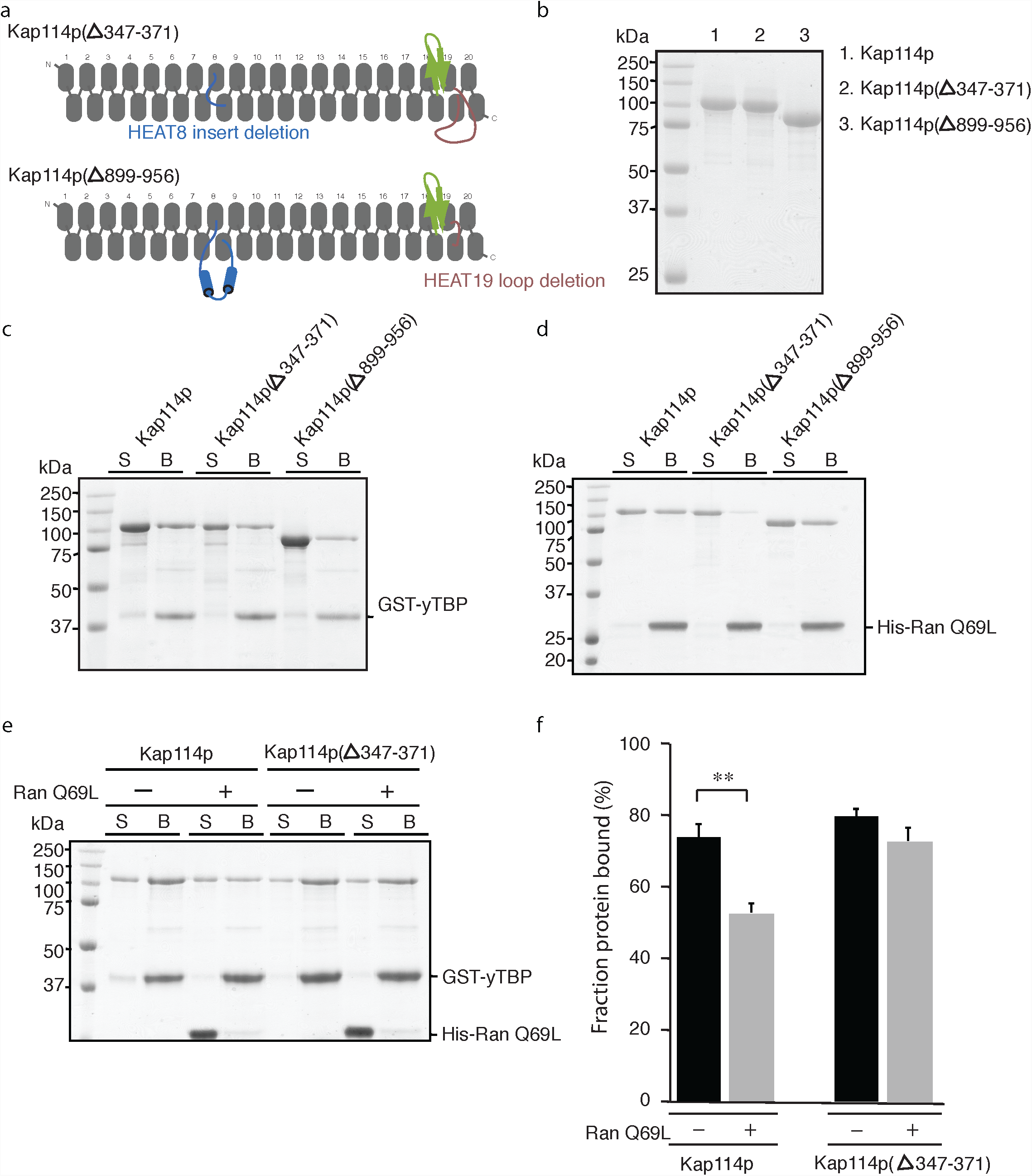
The HEAT8 hairpin of Kap114p is cruical for Ran binding. **a,** Schematic represetation of the domain structures of the two deletion mutants, Kap114p (Δ347-371) and Kap114p (Δ899-956). **b,** Purified recombinant Kap114p, Kap114p (Δ347-371) and Kap114p (Δ899-956) were analyzed by SDS-PAGE, followed by Coomassie blue staining. **c,d,** Pull-down assays of yTBP and RanQ69L (61-240) with Kap114p, Kap114p (Δ347-371) and Kap114p (Δ899-956). **(c)** GST-fused yTBP (61-240) and **(d)** His-tagged RanQ69L were incubated with recombinant Kap114p, Kap114p (Δ347-371) and Kap114p (Δ899-956). Unbound (S) and bound (B) samples were analyzed by SDS-PAGE and stained with Coomassie blue. **e,f,** GST pull-down assays of yTBP (61-240) with Kap114p or Kap114p (Δ347-371) in the presence or absence of RanQ69L. GST-fused yTBP (61-240) was incubated with recombinant Kap114p and His-tagged RanQ69L at a molar ratio of 1:4:8 Kap114p:yTBP:RanQ69L. Unbound (S) and bound (B) samples were analyzed by SDS-PAGE and stained with Coomassie blue. **f,** Analysis of Kap114p dissociation from yTBP in the presence of RanQ69L. Band intensities of Kap114p and Kap114p (Δ347-371) from the SDS– PAGE gels were used to determine the average fraction of bound protein. Data represent mean ± standard deviation from three independent experiments. Differences were assessed statistically by two-tailed Student’s *t* test; **, P < 0.001.

We then performed biochemical pull-down assays using recombinant proteins to assess if the HEAT8 hairpin and HEAT19 loop of Kap114p contribute to the binding of Ran and yTBP. Under our pull-down conditions, both of the HEAT8 hairpin and HEAT19 loop deletion mutants retained the ability to interact with GST-yTBP at levels similar to wild-type Kap114p (Fig. 3c), suggesting that these two segments of Kap114p do not play a major role in yTBP binding. Interestingly, pull-down of Kap114p (Δ347-371) by His-Ran Q69L was substantially diminished compared to that of wild-type and the Kap114p (Δ899-956) mutant (Fig. 3d), suggesting that the HEAT8 hairpin is crucial for Ran binding, as is the case for Importin-β ^16,17^. Furthermore, in a GST pull-down assay with a 1:4:8 mix ratio of Kap114p:yTBP:RanQ69L, wild-type Kap114p and Kap114p (Δ347-371) showed comparable SDS-PAGE band intensities in the protein-bound fractions to assays in the absence of Ran Q69L (Fig. 3e,f), consistent with the fact that they have comparable binding affinities to yTBP. As expected, pull-down of wild type Kap114p by GST-yTBP was significantly different in the presence or absence of Ran Q69L (Fig. 3e,f). However, the amounts of Kap114p (Δ347-371) pulled down by GST-yTBP displayed minimal differences regardless of the presence or absence of Ran Q69L (Fig. 3e,f). Taken together, these results suggest that HEAT8 hairpin-mediated binding of Ran partially dissociates yTBP binding.

### The HEAT8 hairpin of Kap114p suppresses DNA binding by yTBP

Like other TAFs, Kap114p shows a high affinity for yTBP and prevents yTBP homo-dimerization. We sought to establish whether Kap114p has protein features common among TAFs. Interestingly, we found that the HEAT8 hairpin and HEAT19 loop of Kap114p display sequence similarities to the N-terminal domain of TAF1 (TAF1-TAND). TAF1-TAND, which is composed of TAND1 and TAND2 units, modulates interaction of yTBP with TATA box DNA and transcription factors (e.g. TFIIA) ^18,19^. A set of DNA-mimicking residues mediates interaction between two α-helices of TAND1 and the concave surface of yTBP and, remarkably, these residues are also well preserved in the HEAT8 hairpin of Kap114p (Fig. 4a) ^20^. Additionally, the HEAT19 loop of Kap114p is highly negatively charged, presenting protein sequence properties similar to those of TAND2, which represents a conserved regulatory yTBP-binding motif that regulates interaction of yTBP and TFIIA (Fig. 4a) ^20^.

**Fig. 4:**
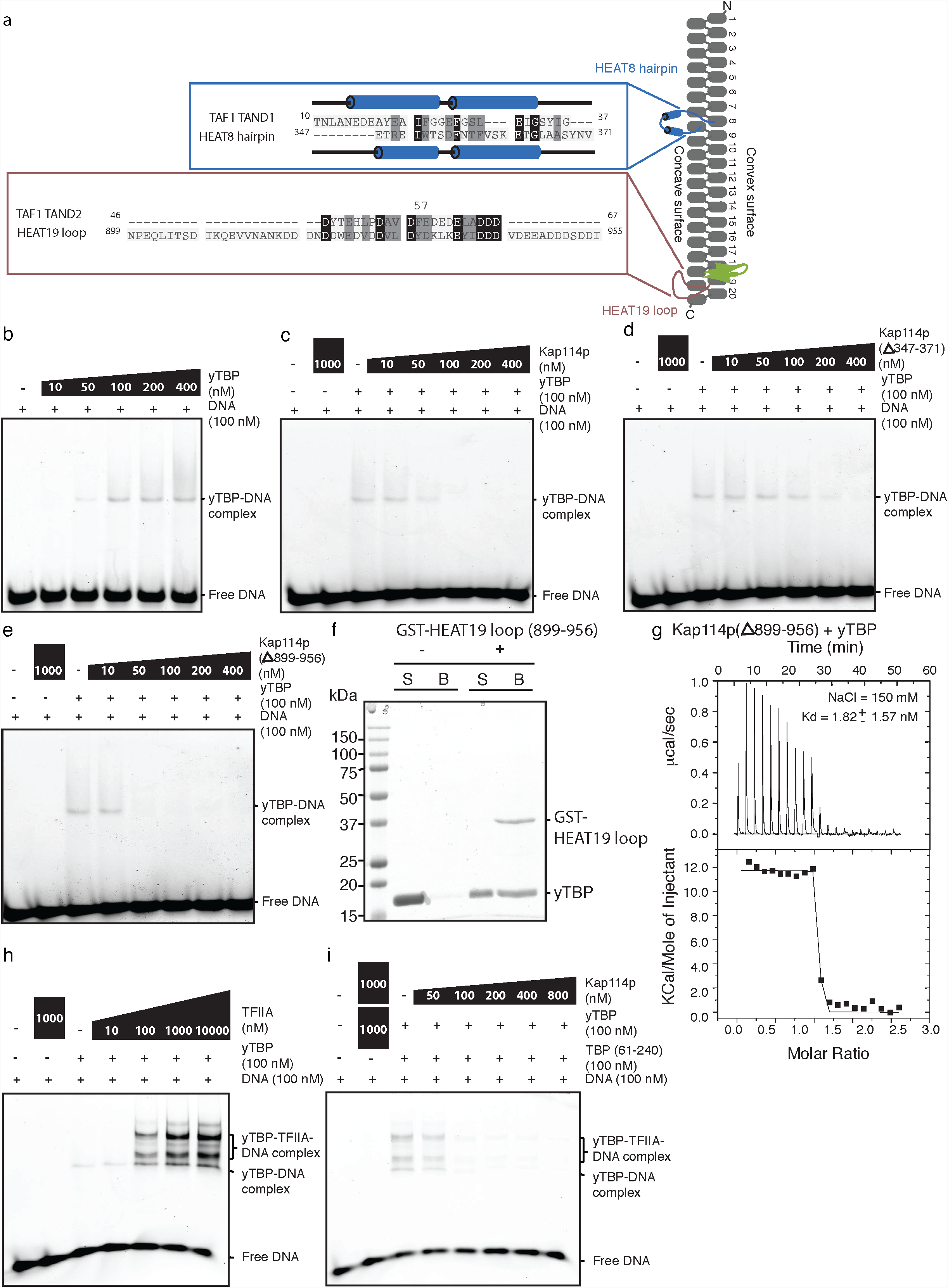
Kap114p modulates yTBP activity. **a,** Protein sequence alignment of the HEAT8 hairpin and TAF1-TAND, and the HEAT19 loop and TAF1-TAND2 (numbers represent amino acid positions). α-helices were assigned based on crystal structures and are depicted as red cylinders. The Phe57 anchor residue of TAND2 is indicated ^20^. **b,** EMSA analysis of yTBP binding to DNA. We incubated 100 nM FAM-labeled oligonucleotide (TGTATGTATATAAAAC) with indicated concentrations of purified yTBP (10 to 400 nM) and then analyzed binding by 6% non-denaturing PAGE. **c,d,e,** Indicated concentrations (10 to 400 nM) of purified recombinant **(c)** Kap114p, **(d)** Kap114p (Δ347-371), and **(e)** Kap114p (Δ899-956) were incubated with yTBP (100 nM) and FAM-labeled DNA (100 nM), followed by native PAGE analysis. WT and Kap114p mutants (1000 nM) were incubated with DNA without yTBP as a control. Electrophoretic bands containing yTBP-DNA complex and free DNA are indicated. **(f)** A GST pull-down assay of the HEAT19 loop (899-956) with yTBP (61-240). yTBP was incubated with and without GST-fused HEAT19 loop (899-956) and pulled down by GST beads. Unbound (S) and bound (B) samples were analyzed by SDS-PAGE and stained with Coomassie blue. **(g)** ITC titration curves (upper) and binding isotherms (lower) of Kap114p (Δ899-956) with yTBP (61-240). Salt concentration and Kd values are indicated. **h,i,** 100nM of TFIIA, yTBP and FAM-labeled DNA (TGTATGTATATAAAAC) were incubated in the **(h)** absence or **(i)** presnece of purified Kap114p, before conducting native PAGE analysis. TFIIA and Kap114p (1000 nM) were incubated with DNA without yTBP as a control. Electrophoretic bands containing protein-DNA complex and free DNA are indicated.

The separation of the HEAT8 hairpin and HEAT19 loop is greater than 500 residues, i.e. vastly different to the 10 residues separating TAND1 and TAND2. We next tested through electrophoretic mobility shift assay (EMSA) whether these two Kap114p inserts could regulate activity of yTBP in a way similar to TAND. Under our experimental conditions, yTBP was able to bind the FAM (Fluorescein amidite) 5’-labeled TATA-box double-stranded DNA in a concentration-dependent manner, leading to band shift to a higher position on the gel (Fig. 4b). Interestingly, the shifted bands induced by yTBP (100 nM) disappeared when more than 100 nM of Kap114p was used (Fig. 4c), suggesting that Kap114p prevents interaction of yTBP and DNA. Furthermore, though wild-type Kap114p and Kap114p (Δ899-956) showed comparable efficiencies in blocking the DNA binding of yTBP, we had to add up to 400 nM of Kap114p (Δ347-371) before the band shift no longer occurred (Fig. 4d,e). Thus, the HEAT8 hairpin of Kap114p plays an important role in suppressing DNA-binding by yTBP.

### The HEAT19 loop of Kap114p interacts with TFIIA

The HEAT19 loop contributes minimally to the binding affinity of Kap114p and yTBP, as the Kd value between Kap114p (Δ899-956) and yTBP is comparable to that of wild-type Kap114p, as determined by ITC (Fig. 1c and 4g). However, disappearance of the biphasic binding profile revealed by ITC analysis suggests that the HEAT19 loop does contact yTBP. We confirmed this weak contact by GST pull-down analysis, evidenced by GST-fused HEAT19 loop (a.a. 899-956) pulling down yTBP under low stringency conditions (50 mM NaCl) (Fig. 4f). Furthermore, we carried out super-shift EMSA using recombinant TFIIA to examine whether Kap114p blocks yTBP and TFIIA interaction ^21,22^. Under our experimental conditions, TFIIA induced super-shift bands on the gel in a dose-dependent manner in the presence of yTBP (Fig. 4h), consistent with TFIIA being able to stabilize yTBP and DNA interaction to form a ternary complex. Moreover, the intensities of the super-shifted bands that represent the ternary complex were dramatically reduced when we applied increasing concentrations of wild-type Kap114p to the reaction (Fig. 4i), indicating suppression of ternary complex formation by Kap114p. Notably, under the same acquisition conditions, the super-shifted bands were still observed when up to 800 nM of either Kap114p (Δ347-371) or Kap114p (Δ899-956) was used compared to when 50 nM of wild-type Kap114p was employed (Supplementary Fig. 3f,g). Thus, deletion of either the HEAT8 hairpin or the HEAT19 loop is insufficient to inhibit formation of the ternary complex since its assembly is facilitated by multiple protein-protein interactions (Fig. 5c). Our analyses indicate that the three-dimensional arrangement of Kap114p allows the HEAT8 hairpin and HEAT19 loop to work in concert, performing a function similar to that of TAF1-TAND.

**Fig. 5:**
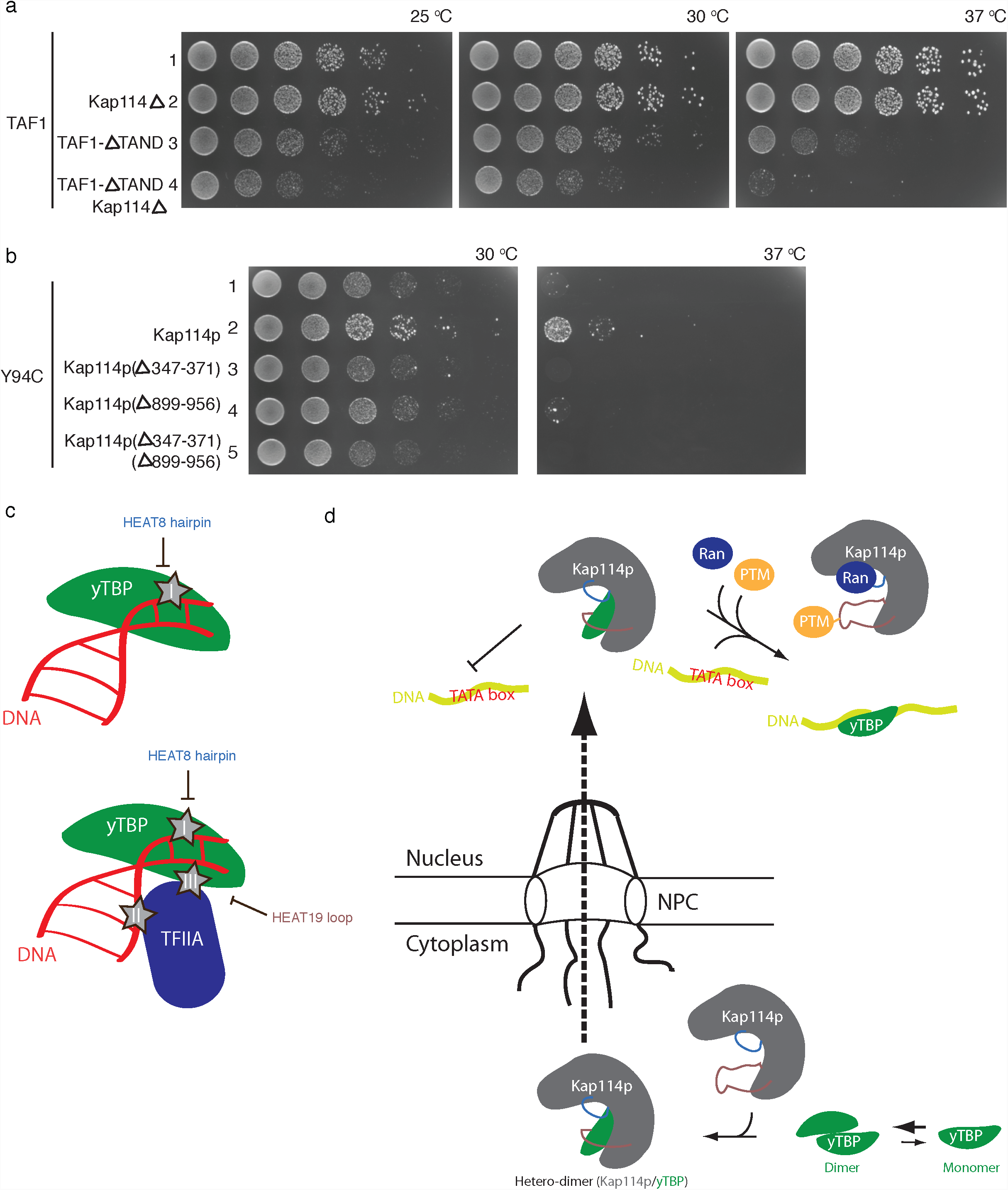
Kap114 and TAF1 work synergistically. **a.** The yTBP temperature-sensitive mutant (Y94C) was transformed with vector control (lane 1) and a plasmid containing wild-type *Kap114, Kap114 (Δ347-371), Kap114 (Δ899-956)* or *Kap114p (Δ347-371, Δ899-956)* (lanes 2-5). **b,** *Kap114* was knocked out in strains bearing wild-type *TAF1* (lane 2) or *ΔTAF1-TAND* (lane 4). Growth effects are shown following serial-dilution (five-fold) of strains. Growth temperatures are indicated. **c,** Schematic representation of the yTBP.DNA and TFIIA.yTBP.DNA complexes. Major contacts within the ternary complex are indicated by gray stars. Contact regions proposed to be blocked by the HEAT8 hairpin and HEAT19 loop are indicated. **d,** Schematic representation of nuclear transport and yTBP activity regulation.

### Synergistic effects of *Kap114* and *TAF1*

Next, we applied yeast genetic approaches to examine our biochemical findings of TAF1-TAND-like motifs in Kap114p in a cellular context. Notably, although knockout of *Kap114* in yeast did not result in any growth defect (Fig. 5a lanes 1 and 2), dual-knockout of *Kap114* and *TAF1-TAND1* caused reduced cell growth when compared to knockout of *TAF1-TAND1* alone (*ΔTAF1-TAND1*) (Fig. 5a lanes 3 and 4) 20. Since the double-knockout mutant elicits a stronger effect than the additive effects of individual mutants, *Kap114* and *TAF1* must be functionally related genes and have synergistic effects on the function of yTBP.

Furthermore, expression of wild-type Kap114p (driven by an endogenous promoter) improved cell growth in a temperature-sensitive yTBP mutant (Y94C) at 37 ^o^C demonstrated by yeast spot-based assays, but the HEAT8 hairpin-and HEAT19 loop-deletion mutants did not exhibit the same effect (Fig. 5b). These results suggest that the HEAT8 hairpin and HEAT19 loop play an important role in complementing the temperature sensitivity caused by this particular yTBP mutation (Y94C), and this activity is unlikely via nuclear transport ^7^. Taken together, these yeast genetics results reinforce our biochemical findings that Kap114p has a dual function and modulates yTBP activity via TAF1-TAND-like motifs.

## DISCUSSSION

Gene duplication and functional divergence have been proposed as important evolutionary mechanisms driving expansion of the Karyopherin-β protein family in order to endow it with the diverse functions that it carries out ^23^. Multiple lines of evidence suggest that a paralogous gene pair, probably created by gene duplication, encodes Kap114p and Cse1p. First, among Kap-βs, Kap114p has been shown to be the most similar to Cse1p at the level of primary protein sequence (18% identity and 43% similarity) ^7^. Second, our crystal structure of Kap114p further revealed high structural similarity to Cse1p, including two HEAT repeat inserts (HEAT8 hairpin and HEAT19 loop). Third, both the *Kap114* and *Cse1* genes are located on chromosome VII and are separated by only ∼ 4,000 nucleotides. Functional divergence may occur after gene duplication, resulting in a common backbone with largely different electrostatic potentials on the concave surface, as well as different properties of hosted intra-HEAT repeat inserts, both of which are important for accommodating the nuclear import of highly basic proteins.

Multiple Kap-βs have been shown to mediate nuclear import of yTBP (e.g. Kap95p, Kap114p, Kap121p and Kap123p), but temperature-sensitive phenotypes caused by yTBP mutations were specifically suppressed by Kap114p ^7^. Interestingly, these mutations (Y94C, K133 and 138L) localize to the DNA-binding pocket and TFIIA interaction site of yTBP ^7,21,24^. Hence, in addition to nuclear import, we propose that Kap114p may carry out other functions that facilitate rescue of the yTBP mutant phenotypes. This hypothesis is supported by our own observations, presented herein, and evidence from other studies. Firstly, knockdown of Kap114p in yeast only partially perturbed yTBP nuclear localization ^7,8^, suggesting that Kap114p is not an essential transport factor for yTBP. Secondly, the Kd values for Kap114p and other Kap-βs we report here suggest that yTBP preferably binds to Kap114p relative to other Kap-βs under cellular conditions, since concentrations of Kap95p, Kap121p and Kap114p in yeast are comparable ^25^. Thirdly, Kap114p displays a stronger affinity for yTBP but a weaker affinity for Ran compared to other Kap-βs ^25^, suggesting a low efficiency for discharging cargoes in the nucleus. Finally, Imp9 (a human homolog of Kap114p) has been proposed to have the dual function of nuclear import factor and cytoplasmic chaperone preventing aggregation of highly basic proteins ^26^.

TAF1 interacts with yTBP via its TAND component comprising TAND1 and TAND2 to modulate gene transcription. Hydrophobic residues on two α-helices of TAND1 that mimic base or ribose moieties of DNA facilitate DNA interaction ^20^. Negatively-charged residues of TAND2, together with an aromatic amino acid, interact with highly basic yTBP ^20^. TAND2 represents a conserved and aspartic acid/glutamic acid-rich TBP-binding motif that has been identified in many structurally distinct proteins such as TFIIA, Brf1 and Mot1 ^27^. Notably, the HEAT8 hairpin and HEAT19 loop of Kap114p display protein sequence compositions and secondary structure elements similar to TAND1 and TAND2, respectively. Our biochemical analyses further reveal that the HEAT8 hairpin and HEAT19 loop regulate yTBP activity by suppressing DNA and TFIIA interactions, respectively. These two intra-HEAT repeat inserts are ∼ 500 residues apart, making it difficult for them to be identified by traditional bioinformatics approaches. However, our structural and biochemical results suggest that the three-dimensional arrangement of Kap114p allows these two HEAT-repeat inserts to operate in concert, thereby conducting roles comparable to those performed by TAF1-TAND. Hence, Kap114p not only acts a transport factor mediating yTBP nuclear import, but it also functions as a TBF-associated factor that negatively regulates yTBP activity.

Unlike for other Kap-βs, Ran-GTP only partially disassembles Kap114p.yTBP complexes, so additional mechanisms that unload yTBP in the nucleus have been proposed ^8^. For instance, TATA box-containing DNA has been shown to enhance dissociation of yTBP and Kap114p in the presence of Ran ^8^. Here, we have biochemically demonstrated that deletion of the HEAT8 hairpin not only reduced the affinity of Kap114p for Ran, but also inhibited yTBP and DNA interaction. Accordingly, we propose a model of the molecular mechanism underlying DNA-enhanced cargo dissociation by Kap114p whereby Ran and yTBP compete for binding to the HEAT8 hairpin of Kap114p. Binding of Ran with the HEAT8 hairpin may expose the concave DNA-binding surface of yTBP originally occluded by the HEAT8 hairpin, allowing the DNA to further promote dissociation yTBP and Kap114p. Moreover, post-translational modification by sumolyation of Kap114p residue 909 has been proposed to release cargoes from Kap114p ^28^. Lysine residue 909 lies within the unstructured HEAT19 loop according to our Kap114p crystal structure. Since our biochemical analyses suggest that the HEAT19 loop does not contribute significantly to yTBP binding but instead interferes with TFIIA interaction, it is possible that sumolyation releases the contact between the HEAT19 loop and yTBP, working together with DNA and Ran to fully discharge yTPB (Fig. 5d).

Nuclear transport and competitive inhibition of yTBP were originally deemed to be conducted by two separate groups of proteins (i.e. nuclear transport factors and TBP-associated factors). However, coupling of Kap114p nuclear import to yTBP nuclear activity regulation allows cells to efficiently tune gene expression in response to environmental change. Furthermore, the high-affinity binding between Kap114p and yTBP that is regulated by multiple factors (Ran, DNA and post-translational modification) precisely controls yTBP activity in the nucleus (Fig. 5d). Hence, the combinatorial regulation conducted by Kap114p is an efficient way of precisely controlling gene expression.

## MATERIALS AND METHODS

### Protein expression and purification

Yeast (*S. cerevisiae*) full-length Kap95p, Kap114p, and Kap121p were amplified by PCR and inserted into the pGEX6p1 expression vector (GE Healthcare) that contains a PreScission cleavage site. Proteins were expressed using the *Escherichia coli* Rosetta strain (Novagen). Cell culture was induced by overnight incubation with 0.5 mM IPTG at 18 °C. Cells were harvested and resuspended in lysis buffer [20 mM HEPES (pH 7.4), 150 mM NaCl, 3 mM DTT]. The cells were lysed by French press and the lysate was centrifuged for 30 min at 15,000 *g*. The supernatant was incubated with GST resins (GE Healthcare), and the GST-fused protein was then eluted by a buffer containing 50 mM of reduced glutathione. The GST-tag was removed by PreScission protease before protein samples were further purified through HiTrap Q HP columns and SEC (Superdex 200 16/60). Protein quality was analyzed and confirmed by SDS-PAGE. Deletion mutants of Kap114p were expressed and purified using an identical protocol.

Yeast (*S. cerevisiae*) yTBP (61-240) was cloned into a modified pET28a vector with a PreScission cleavage site at the N-terminus. The same bacterial strain and protein overexpression strategy used to express Kap proteins was applied to express His-yTBP. Cells were harvested and suspended in phosphate buffer containing 150 mM NaCl and 3 mM β-mercaptoethanol. After cell lysis, the lysate was clarified by centrifugation and the supernatant was incubated with Ni beads (Qiagen) for 30 mins, followed by a prewash with 25 mM imidazole. Proteins were eluted with buffer containing 250 mM imidazole, and the fractions were pooled and dialyzed against buffer [20 mM HEPES (pH 7.4), 150 mM NaCl, 3 mM DTT] overnight. After dialysis, protein samples were loaded into HiTrap SP HP columns and eluted by a salt gradient. Protein samples were then further purified by SEC (Superdex 200 16/60) and analyzed by SDS-PAGE. TFIIA was expressed and purified using an established protocol ^22^. All proteins were concentrated and stored at −80 °C. All constructs are listed in Supplementary Table 3.

### Protein crystallization and structure determination

Native Kap114p crystals were obtained using a hanging drop vapor diffusion method with a 17.6 mg/ml protein concentration at 20 °C by mixing 1 ml protein with 1 ml reservoir solution of 0.43 M citric acid, 0.057 M BIS TRIS propane (pH 5.8), 17% PEG3350, 0.01 M Ethylenediaminetetraacetic acid disodium salt dihydrate, and 3% v/v ethanol. Selenium-labeled crystals were grown in 0.044 M citric acid, 0.056 M BIS TRIS propane (pH 5.6), 15% PEG3350, and 0.5% w/v polyvinylpyrrolidone. Crystals appeared in about one week. Crystals were hexagonal (space group P3_1_2_1_), with one molecule per asymmetric unit. Data were collected at beamline 13B, 13C and 05A in the National Synchotron Radiation Research Centre (Taiwan). X-ray intensities were processed using HKL2000 ^29^. Phase determination was carried out using data collected from seleno-L-methionine-labeled crystals at selenium peak and inflection wavelengths using PHENIX ^30^. The initial model was built into the electron density map and refined using PHENIX and Coot ^30,31^. The final structure was refined to a resolution of 2.5 Å with an R-factor of 20.6% (Rfree 25.5%), and there were no outliers in the Ramachandran plot.

### Pull-down assays

GST-tagged yTBP (61-240) (1 mM) was preincubated with MagnetGST^™^ Glutathione Particles (Promega) in binding buffer containing 20 mM HEPES (pH 7.4), 150 mM NaCl, and 3 mM DTT, on ice for 30 mins, before adding 1 mM of Kap114p, Kap114p (Δ347-371) or Kap114p (Δ899-956) for incubation for another 30 mins. The reactions were washed three times using binding buffer and then eluted by 50 mM glutathione. The eluted samples were analyzed by SDS-PAGE. His-tagged RanQ69L (4 mM) was preincubated on ice with MagnetHis^™^ Ni-Particles (Promega) in phosphate binding buffer with 150 mM NaCl and 3 mM β-mercaptoethanol for 30 mins. We added 1 mM of Kap95p, Kap114p, or Kap121p to the reaction before incubating it for another 30 mins on ice. The reactions were washed three times and eluted by buffer containing 250 mM imidazole, and analyzed by SDS-PAGE. For competitive binding analysis (Fig. 4e,f), Kap114p (0.5 mM), GST-tagged yTBP (61-240) (2 mM) and His-tagged RanQ69L (4 mM) were preincubated and pulled down by MagnetGST^™^ Glutathione Particles (Promega).

### Small Angle X-ray Scattering

Purified Kap114, Kap114.RanQ60L, and Kap114p.yTBP complexes were diluted to 5 mg/ml in buffer [20 mM HEPES (pH 7.4), 150 mM NaCl, 3 mM DTT] before being subjected to SAXS analysis. Data were collected at the SAXS beamline 23A in the Taiwan Light Source of the National Synchrotron Radiation Research Centre ^32,33^. The experimental setup consists of a High Pressure Liquid Chromatography (HPLC, Agilent chromatographic system 1260 series) system equipped with an Agilent silica-based column of pore size 300 Å, followed downstream by a SAXS sample capillary. An X-ray beam with wavelength of 0.8266 Å was used for data collection. Selected frames were merged and analyzed for initial *R*_g_ estimation by the PRIMUS program, and the *P*(*r*) distance distribution was determined by the GNOM program, both from the ATSAS package ^34^. Low-resolution *ab initio* envelopes were calculated using DAMMIF and GASBOR. DAMAVER was then used to generate an average model and the crystal structures were fitted into the envelope by SUPCOMB. SAXS modeling of the Kap114p.Ran and Kap114p.yTBP complexes was performed using FoXSDock ^35,36^.

### Isothermal Titration Calorimetry

Binding affinities between yTBP and Kap114p, Kap114p (Δ899-956), Kap95p, and Kap121p were measured by ITC (MicroCal iTC200). All proteins were dialyzed against ITC buffer [20 mM HEPES (pH 7.4), 150 mM NaCl, 1 mM β-mercaptoethanol]. Different Kap proteins were stored in the sample cell and yTBP was injected into the cell by syringe. ITC was performed at 25 °C.

### Electrophoretic mobility shift assay

Electrophoretic mobility shift assay (EMSA) was performed using purified recombinant yTBP, TFIIA, Kap95p, Kap121p, Kap114p, Kap114p (Δ347-371), and Kap114p (Δ899-956). We used synthesized FAM (Fluorescein amidite) 5’-labeled TATA-box double-stranded DNA (TGTATGTATATAAAAC). Samples were analyzed by 4.5% polyacrylamide gels [4.5% acrylamide from a 29%:1% acrylamide:bisacrylamide stock, 25 mM Tris-HCl (pH 8.3), 190 mM glycine, 10 mM EDTA] using TGE running buffer [25 mM Tris-HCl (pH 8.3), 190 mM glycine, 10 mM EDTA]. Gels were pre-run at 100 V at 4 °C for 1 hour before sample loading. Proteins and DNA were incubated on ice at 60 minutes in 20 mM HEPES, 100 mM NaCl, 3 mM DTT, 8% glycerol, and 4 mg bovine serum albumin (BSA).

### Analytic ultracentrifugation

Sedimentation velocity analyses were conducted at 45,000 rpm at 20 °C with a Beckman Optima XL-I AUC system equipped with absorbance optics. Purified protein samples of yTBP, Kap114p, and Kap114p.yTBP were diluted to final concentrations of 1.1 mg/ml, 0.86 mg/ml, and 1.47 mg/ml, respectively. The dilution was performed using 20 mM HEPES (pH 7.4), 150 mM NaCl, 1 mM DTT prior to analysis. Standard 12 mm aluminum double-sector centerpieces were filled with the protein solution, whereas blank buffer was used in the reference cell. Before each run, cells were thermally equilibrated for at least 1 hour in the 4-hole (AnTi60) rotor of the instrument. Quartz windows were used with absorbance optics (OD 280 nm) in continuous mode without averaging. No time interval was set between scans. For sedimentation equilibrium experiments, the Kap114p.yTBP complex was diluted to 0.84 mg/ml and analyzed at 6600, 7900, and 14 000 rpm using a 4-hole AnTi60 rotor at 20 °C. Data were analyzed with a c(s) distribution of Lamm equation solutions calculated by the program SEDFIT (www.analyticalultracentrifugation.com), assuming the regularization parameter p to be 0.95 (high confidence level). The weighted average sedimentation coefficient (S) was obtained by integration over the range of each peak. All S values were corrected for the viscosity and density of water at 20 °C.

### Yeast strain construction

*Kap114* (including the upstream 500 base pairs (bp) and downstream 100 bp) was cloned into the vector pRS426. To examine whether Kap114p deletion mutants rescue the phenotype of the temperature-sensitive yTBP (Y94C) mutation, yeast strain YSB66 ^7^ was transformed with pRS426 as a control or pRS426 carrying wild-type *Kap114, Kap114 (Δ347-371), Kap114 (Δ899-956)*, or *Kap114 (Δ347-371, Δ899-956)*. To test the effect of *Kap114* knockout in the absence of *TAF1-TAND*, we deleted *KAP114* from the YTK12029 and YTK12803 strains ^20^ by replacing it with a gene encoding LEU2. We amplified the LEU2 sequence from pFA6a LEU2 template using the forward primer AAAATCTTGAACGTAATTGTAACACTATCAACACATTAAACGGATCCCCGGGTTAATTAA and the reverse primer CTACTTTACATCTGATATCTCCACGGCTTATGTATATAAGGAATTCGAGCTCGTTTAAAC. The PCR product was transformed into both strains. Genomic DNA of colonies grown on the selective medium was isolated and verified by PCR reaction (Supplementary Fig. 3h,i,j). All yeast strains used in this study are listed in Supplementary Table 2.

### Yeast spot-based assay

For dilution spot assays, yeast samples grown to logarithmic phase were initially diluted to the same OD. Subsequently, a series of five-fold dilutions was generated, and then 10 μl of each sample was spotted onto selective medium and incubated at the indicated temperatures.

## REFERENCES

1 Vannini, A. & Cramer, P. Conservation between the RNA polymerase I, II, and III transcription initiation machineries. Mol Cell 45, 439–46 (2012).

2 Hahn, S. Structure and mechanism of the RNA polymerase II transcription machinery. Nat Struct Mol Biol 11, 394–403 (2004).

3 Coleman, R.A., Taggart, A.K., Benjamin, L.R. & Pugh, B.F. Dimerization of the TATA binding protein. J Biol Chem 270, 13842–9 (1995).

4 Chasman, D.I., Flaherty, K.M., Sharp, P.A. & Kornberg, R.D. Crystal structure of yeast TATA-binding protein and model for interaction with DNA. Proc Natl Acad Sci U S A 90, 8174–8 (1993).

5 Nikolov, D.B. et al. Crystal structure of TFIID TATA-box binding protein. Nature 360, 40–6 (1992).

6 Chitikila, C., Huisinga, K.L., Irvin, J.D., Basehoar, A.D. & Pugh, B.F. Interplay of TBP inhibitors in global transcriptional control. Mol Cell 10, 871–82 (2002).

7 Morehouse, H., Buratowski, R.M., Silver, P.A. & Buratowski, S. The importin/karyopherin Kap114 mediates the nuclear import of TATA-binding protein. Proc Natl Acad Sci U S A 96, 12542–7 (1999).

8 Pemberton, L.F., Rosenblum, J.S. & Blobel, G. Nuclear import of the TATA-binding protein: mediation by the karyopherin Kap114p and a possible mechanism for intranuclear targeting. J Cell Biol 145, 1407–17 (1999).

9 Matsuura, Y. & Stewart, M. Structural basis for the assembly of a nuclear export complex. Nature 432, 872–7 (2004).

10 Kutay, U., Bischoff, F.R., Kostka, S., Kraft, R. & Gorlich, D. Export of importin alpha from the nucleus is mediated by a specific nuclear transport factor. Cell 90, 1061–71 (1997).

11 Matsuura, Y. Mechanistic Insights from Structural Analyses of Ran-GTPase-Driven Nuclear Export of Proteins and RNAs. J Mol Biol 428, 2025–39 (2016).

12 Greiner, M., Caesar, S. & Schlenstedt, G. The histones H2A/H2B and H3/H4 are imported into the yeast nucleus by different mechanisms. Eur J Cell Biol 83, 511–20 (2004).

13 Mosammaparast, N., Ewart, C.S. & Pemberton, L.F. A role for nucleosome assembly protein 1 in the nuclear transport of histones H2A and H2B. EMBO J 21, 6527–38 (2002).

14 Fox, A.M., Ciziene, D., McLaughlin, S.H. & Stewart, M. Electrostatic interactions involving the extreme C terminus of nuclear export factor CRM1 modulate its affinity for cargo. J Biol Chem 286, 29325–35 (2011).

15 Christie, M. et al. Structural Biology and Regulation of Protein Import into the Nucleus. J Mol Biol 428, 2060–90 (2016).

16 Conti, E. & Izaurralde, E. Nucleocytoplasmic transport enters the atomic age. Curr Opin Cell Biol 13, 310–9 (2001).

17 Chook, Y.M. & Blobel, G. Karyopherins and nuclear import. Curr Opin Struct Biol 11, 703–15 (2001).

18 Mal, T.K. et al. Structural and functional characterization on the interaction of yeast TFIID subunit TAF1 with TATA-binding protein. J Mol Biol 339, 681–93 (2004).

19 Bagby, S. et al. TFIIA-TAF regulatory interplay: NMR evidence for overlapping binding sites on TBP. FEBS Lett 468, 149–54 (2000).

20 Anandapadamanaban, M. et al. High-resolution structure of TBP with TAF1 reveals anchoring patterns in transcriptional regulation. Nat Struct Mol Biol 20, 1008–14 (2013).

21 Tan, S., Hunziker, Y., Sargent, D.F. & Richmond, T.J. Crystal structure of a yeast TFIIA/TBP/DNA complex. Nature 381, 127–51 (1996).

22 Adachi, N. et al. Improved method for soluble expression and rapid purification of yeast TFIIA. Protein Expr Purif 133, 50–56 (2017).

23 O’Reilly, A.J., Dacks, J.B. & Field, M.C. Evolution of the karyopherin-beta family of nucleocytoplasmic transport factors; ancient origins and continued specialization. PLoS One 6, e19308 (2011).

24 Kim, Y., Geiger, J.H., Hahn, S. & Sigler, P.B. Crystal structure of a yeast TBP/TATA-box complex. Nature 365, 512–20 (1993).

25 Hahn, S. & Schlenstedt, G. Importin beta-type nuclear transport receptors have distinct binding affinities for Ran-GTP. Biochem Biophys Res Commun 406, 383–8 (2011).

26 Jakel, S. et al. The importin beta/importin 7 heterodimer is a functional nuclear import receptor for histone H1. EMBO J 18, 2411–23 (1999).

27 Chou, C.C. & Wang, A.H. Structural D/E-rich repeats play multiple roles especially in gene regulation through DNA/RNA mimicry. Mol Biosyst 11, 2144–51 (2015).

28 Rothenbusch, U., Sawatzki, M., Chang, Y., Caesar, S. & Schlenstedt, G. Sumoylation regulates Kap114-mediated nuclear transport. EMBO J 31, 2461–72 (2012).

29 Otwinowski, Z. & Minor, W. Processing of X-ray diffraction data collected in oscillation mode. Methods Enzymol 276, 307–26 (1997).

30 Adams, P.D. et al. PHENIX: a comprehensive Python-based system for macromolecular structure solution. Acta Crystallogr D Biol Crystallogr 66, 213–21 (2010).

31 Emsley, P. & Cowtan, K. Coot: model-building tools for molecular graphics. Acta Crystallogr D Biol Crystallogr 60, 2126–32 (2004).

32 Yeh, Y.Q. et al. Probing the Acid-Induced Packing Structure Changes of the Molten Globule Domains of a Protein near Equilibrium Unfolding. J Phys Chem Lett 8, 470–477 (2017).

33 Shih, O. et al. Membrane Charging and Swelling upon Calcium Adsorption as Revealed by Phospholipid Nanodiscs. J Phys Chem Lett 9, 4287–4293 (2018).

34 Franke, D. et al. ATSAS 2.8: a comprehensive data analysis suite for small-angle scattering from macromolecular solutions. J Appl Crystallogr 50, 1212–1225 (2017).

35 Schneidman-Duhovny, D., Hammel, M. & Sali, A. Macromolecular docking restrained by a small angle X-ray scattering profile. J Struct Biol 173, 461– 71 (2011).

36 Schneidman-Duhovny, D., Hammel, M., Tainer, J.A. & Sali, A. FoXS, FoXSDock and MultiFoXS: Single-state and multi-state structural modeling of proteins and their complexes based on SAXS profiles. Nucleic Acids Res 44, W424–9 (2016).

